# Robust Binding Capability and Occasional Gene Loss of Telomere-Binding Proteins Underlying Telomere Evolution in Nematoda

**DOI:** 10.1101/2024.07.11.603171

**Authors:** Hobum Song, Seonhong Kim, Daisy S. Lim, Hee-Jung Choi, Junho Lee

## Abstract

Telomeres, the nucleoprotein complexes that protect the ends of linear chromosomes, are essential for maintaining the stability of eukaryotic genomes. As telomeres generally consist of repetitive DNA associated with specifically bound proteins, telomeric repeat motifs (TRMs) are thought to be difficult to evolve. However, a recent study identified nematodes with telomeric repeats distinct from the canonical TTAGGC motif. Here, we investigated how telomere repeats could have evolved despite the challenge posed by the specificity of telomere-binding proteins (TBPs) to the telomeric DNA in Nematoda. We performed a phylogenetic analysis and electrophoresis mobility shift assays to assess the binding affinities of two TBPs, which displayed different conservation patterns. Our results revealed that the well-conserved protein CEH-37 exhibits limited specificity, unable to distinguish telomeric repeats found in nematodes except for the TTAGGG motif, while the less conserved POT proteins displayed rigid specificity. These findings suggest that the emergence of novel telomeric repeat motifs correlated with the characteristics and evolutionary outcomes of TBPs in Nematoda. Our study not only revealed the dynamics of telomere evolution but also enhanced the understanding of the evolutionary relationship between proteins and DNAs.

**Lay summary:** Telomeres are chromosomal end structures composed of repetitive DNA where telomere-binding proteins (TBPs) specifically bind. This nucleoprotein complex is crucial for eukaryotes to maintain their genome integrity, making it difficult to evolve. However, a recent study identified nematodes with non-canonical telomeric repeat motifs (TRMs). Through an analysis of TBP conservation in Nematoda and assessments of their binding affinities, we observed that the binding capabilities of TBPs align with their conservation patterns. This correlation is also linked to the emergence of novel TRMs, explaining the evolution progress of TRMs in Nematoda. The evolutionary process discovered in this research is not limited to telomeres, providing further insights into evolutionary relationships between proteins and DNA.

## Introduction

Genome stability in eukaryotes is inherently challenged by the continuous shortening of chromosomal ends and their misrecognition as double-strand breaks (De Lange, 2009; Olovnikov, 1973; Watson, 1972). To address these problems, eukaryotes have nucleoprotein structures at the ends of their linear chromosomes called telomeres. Telomere-binding proteins (TBPs) specifically bind to DNA sequences generally composed of G-rich repeat motifs, forming complexes that cap the chromosomal ends to prevent the activation of the DNA damage response and regulate the length of telomeres, ensuring functional telomeres. Consequently, changes in telomeric repeat motifs (TRMs) can lead to genomic instability by affecting the binding affinities of TBPs, making the evolution of telomeric sequences challenging (Steinberg-Neifach and Lue, 2015).

Despite its challenging nature, telomeric sequence evolution has been observed in several lineages, including yeast, plants, and insects (Červenák et al., 2021; Kuznetsova et al., 2020; Procházková Schrumpfová et al., 2016). Various hypotheses have been proposed to explain how this process was possible, and some of these hypotheses have been experimentally validated in certain organisms (Saint-Leandre and Levine, 2020; Steinberg-Neifach and Lue, 2015). In yeast, the adoption of flexible TBPs to cope with rapid changes in telomeric sequences has been observed, while the coevolution of TRM and TBP has been the principle of telomere evolution in plants (Červenák et al., 2021; Sepsiova et al., 2016; Shakirov et al., 2009). In Animalia, however, no studies have been conducted to explain the relationship between TBPs and the evolution of telomeric sequences.

Nematodes are good model organisms with well-constructed databases and systems for research in diverse fields (Howe et al., 2016; Howe et al., 2017). Within a representative model organism *Caenorhabditis elegans*, research on telomeres has been conducted using these models and enabled the identification of *C. elegans* TBPs. While it has recently confirmed that TEBP proteins, POT proteins, and MRT-1 interact to form telomeric protein complexes, there are other TBPs besides those subunits such as CEH-37, HMG-5, PLP-1, and HRPA-1, making Nematoda an intriguing group for studying telomere evolution (Dietz et al., 2021; Im and Lee, 2003; Im and Lee, 2005; Joeng et al., 2004; Kim et al., 2003; Meier et al., 2009; Raices et al., 2008; Yamamoto et al., 2021; Yu et al., 2022).

Recently, novel TRMs have been discovered within the Nematoda phylum, which includes one of the representative model organisms *Caenorhabditis elegans* (Lim et al., 2023). While it has been well established that the nematode TRM is TTAGGC, some species, such as *C. uteleia*, *Panagrellus redivivus* and diverse isolates in Panagrolaimidae family (e.g. LJ2284, LJ2285), and *Strongyloides ratti*, have been revealed to possess noncanonical TRMs (TTAGGT, TTAGAC, and TTAGGG, respectively) by analysis of whole genome sequence (WGS) data (Figure 1A). Given these findings, we aimed to understand how TRM evolved in Nematoda despite the challenges posed by sequence-specificities of TBPs. Based on our phylogenetic analysis, we focused on CEH-37, a well-conserved TBP, and less conserved POT proteins. The binding specificities of these proteins varied according to their conservation patterns – CEH-37 exhibited flexibility, while the POT proteins displayed relatively rigid specificity. These results indicate that the flexibility or the loss and gain of TBPs may coincide with the emergence of novel TRMs.

**Figure 1.**
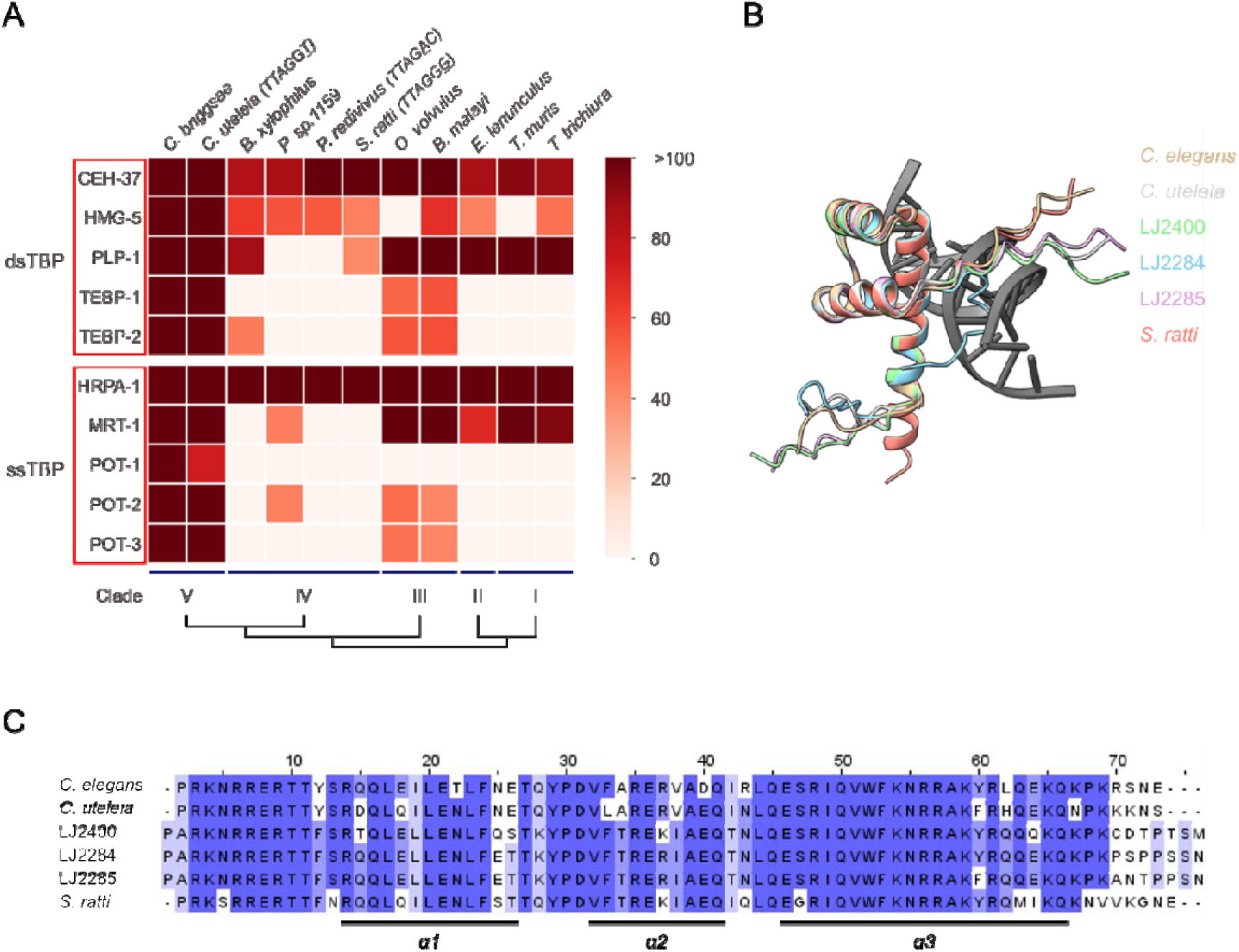
Conservation of telomere binding proteins in nematodes. (A) Heatmap illustrating the conservation of *C. elegans* TBPs in various nematodes according to the bit score. For nematodes with noncanonical TRMs, the TRMs are indicated in brackets after the name of the species. Nematodes with a canonical TTAGGC TRM are not explicitly labeled. Phylogenetic tree under the heatmap is based on small rRNA phylogeny, which serves as the standard for defining nematode clades. (dsTBP: double-stranded telomeric DNA binding protein, ssTBP: single-stranded telomeric DNA binding protein.) (B) Model of the DNA-CEH-37 homeodomain complex of *C. elegans* with CEH-37 homeodomains from other nematodes aligned to it. The color of the species names corresponds to the color of the CEH-37 homeodomains. (C) Multiple sequence alignment result of CEH-37 homeodomains from nematodes used in this study. The brightness of residue colors indicates the percentage identity in each column to the consensus sequence. Each α-helix region is indicated under the result of multiple sequence alignment. Sequence alignment was performed using Clustal Omega.

## Results

### Conservation of *C. elegans* telomere-binding proteins in Nematoda

In eukaryotes, telomeric DNAs are capped by protein complexes (De Lange, 2005; Moser and Nakamura, 2009). In *C. elegans*, several TBPs have been identified, and TEBP-1, TEBP-2, POT-1, POT-2, and MRT-1 are thought to form a telomeric protein complex (Dietz et al., 2021; Meier et al., 2009; Raices et al., 2008; Yamamoto et al., 2021). However, there are some species in Nematoda found to possess noncanonical TRMs (Lim et al., 2023). Since TBPs need to bind effectively to telomeric DNA for telomeres to function properly, we investigated which TBPs of *C. elegans* have been conserved across other nematode species despite differences in their TRMs.

As nematodes are classified into five clades based on their small rRNA phylogeny (Blaxter et al., 1998), we conducted BLAST analysis of TBPs from representative species of each clade, including the ones with noncanonical TRMs (Figure 1A, Supplementary Table S1). Using a bit score threshold of 50 in BLASTP to indicate significant homology (Pearson, 2013), we observed the TBPs of a clade V nematode species *C. elegans* are well conserved within clade V, while it was not the same in other clades. Notably, CEH-37 and HRPA-1 are well conserved across all clades of Nematoda. In contrast, the major subunits of the *C. elegans* telomeric protein complex, TEBP and POT proteins, are not conserved in clade IV, which contains species with the TTAGAC TRM, as well as clades I and II. This suggests that the species in clades other than III and V must possess a telomeric protein complex distinct from that of *C. elegans*. Our analysis shows that TBPs experienced different selection pressure during the evolution and speciation of nematodes.

### Conserved homeodomain of CEH-37 robustly binds to telomeric DNAs with limited binding specificity

CEH-37, the only well-conserved double-stranded telomeric DNA binding protein (dsTBP) in Nematoda, was initially proposed as an evolutionarily conserved TBP due to its structural similarity to dsTBPs in other species, which typically use Myb-related domains as DNA-binding domains (DBDs) (Kim et al., 2003; Moon et al., 2014). However, the discovery of TEBP proteins, which also possess Myb-related domains, raised doubts about this hypothesis. Notably, we observed that certain clades lack TEBP proteins and other subunits of the *C. elegans* telomeric protein complex. To investigate which TBPs might function like representative dsTBPs in other species, we examined structural homology between human TRF1 and *C. elegans* TBPs other than TEBP proteins, using TM-align (Zhang and Skolnick, 2005). Given that a TM-score above 0.5 supports an evolutionary relationship, CEH-37 was the only dsTBP with a DBD structurally similar to the DBD of TRF1 (Supplementary Figure S1, Supplementary Table S2). In addition, the CEH-37 homeodomain displayed structural similarity with other DBDs of representative dsTBPs such as *Schizosaccharomyces pombe* Taz1 and *Saccharomyces cerevisiae* RAP1 (Supplementary Table S2). These unique characteristics of CEH-37 as a TBP raise questions about its role in the evolution of nematode telomeres.

To investigate how novel telomeric sequences emerged despite the conservation of CEH-37, we obtained the recombinant CEH-37 homeodomains from nematodes with different TRMs (Supplementary Figure S2) and measured their binding affinities to telomeric DNAs identified in Nematoda. First, we compared the binding affinities of CEH-37 homeodomains from *C. elegans* and *C. uteleia*, which have canonical TRM TTAGGC and noncanonical TRM TTAGGT, respectively, using electrophoresis mobility shift assay (EMSA) (Figure 2, Supplementary Table S3 and S4). Interestingly, CEH-37 homeodomains from both species bound to telomeric DNAs with TTAGGC, TTAGAC, and TTAGGT repeats, with no significant difference in their dissociation constants (K_d_). However, both CEH-37 homeodomains did not effectively bind to DNA with TTAGGG repeats. These results indicate that CEH-37 does not distinguish TTAGGC, TTAGAC, and TTAGGT motifs, while it shows less preference for TTAGGG motif.

**Figure 2.**
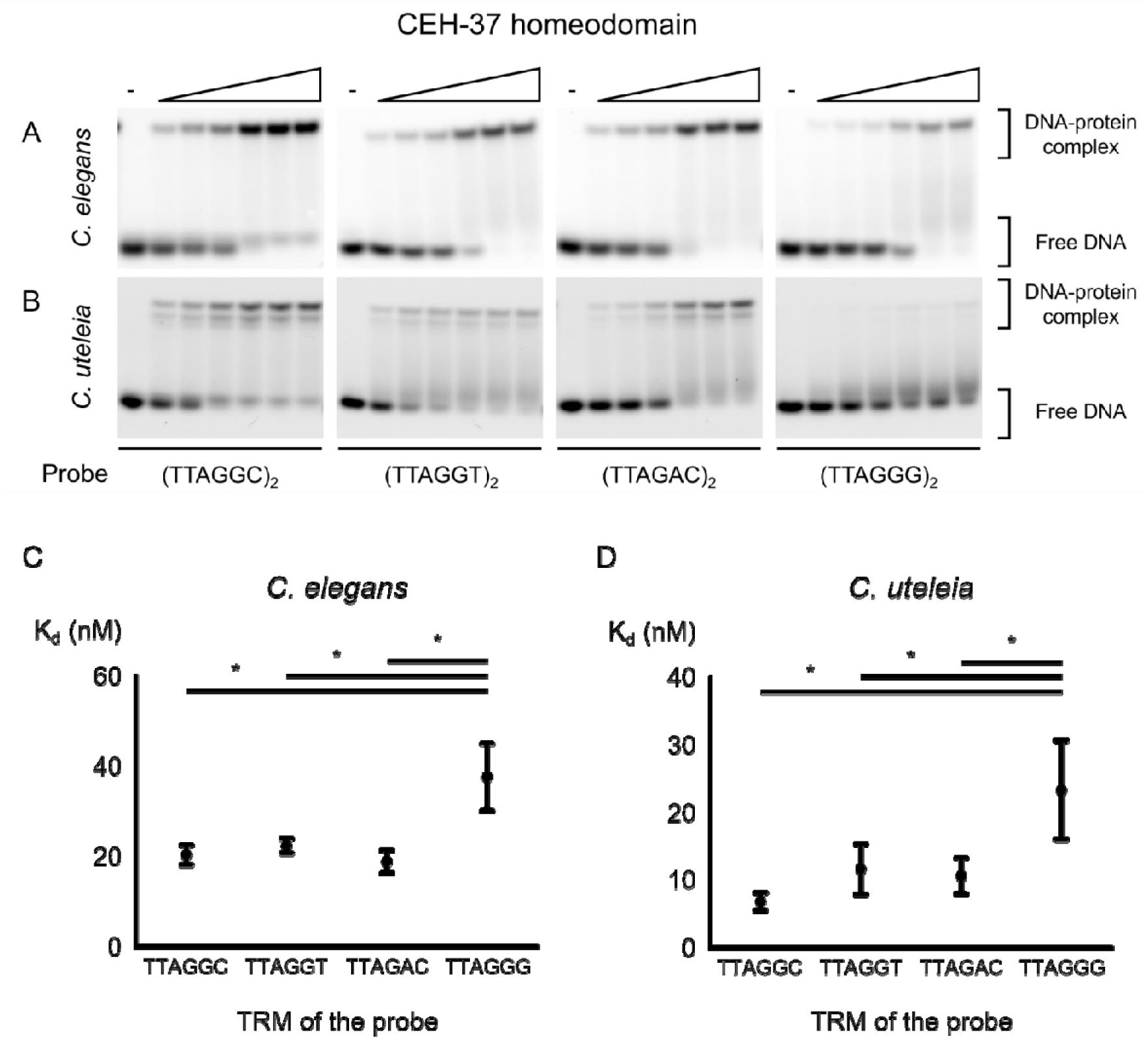
Binding affinity of CEH-37 homeodomains from the *Caenorhabditis* genus. (A, B) Representative images of EMSA demonstrating the binding of the *C. elegans* and *C. uteleia* CEH-37 homeodomains to telomeric DNAs, respectively. 5 nM of DNA probes indicated under each figure were incubated with increasing concentrations of proteins ranging from 5 nM to 160 nM. (C, D) K_d_ of *C. elegans* and *C. uteleia* CEH-37 homeodomains to each telomeric DNA, respectively. DNA probes used for the K_d_ measurements consist of 2 repeats of the indicated sequences. Relationships without asterisks indicate non-significant differences (Newman-Keuls multiple comparison test, * *p*<0.05).

We also measured the binding affinities of CEH-37 homeodomains from strains in clade IV, where species with other noncanonical TRMs were identified (Figure 3, Supplementary Table S3 and S4). The Panagrolaimidae family in clade IV includes isolate LJ2284, which possess noncanonical TRM TTAGAC, and a related species LJ2400, which has canonical TRM TTAGGC. Measurement of K_d_ values of CEH-37 homeodomains from LJ2284 and LJ2400 showed consistent results with the above experiments, indicating robust binding to telomeric DNA with limited sequence specificity, irrespective of the TRM of their origin species. These results suggest that TRMs in Nematoda evolved within the range of the well-conserved CEH-37 sequence specificity.

**Figure 3.**
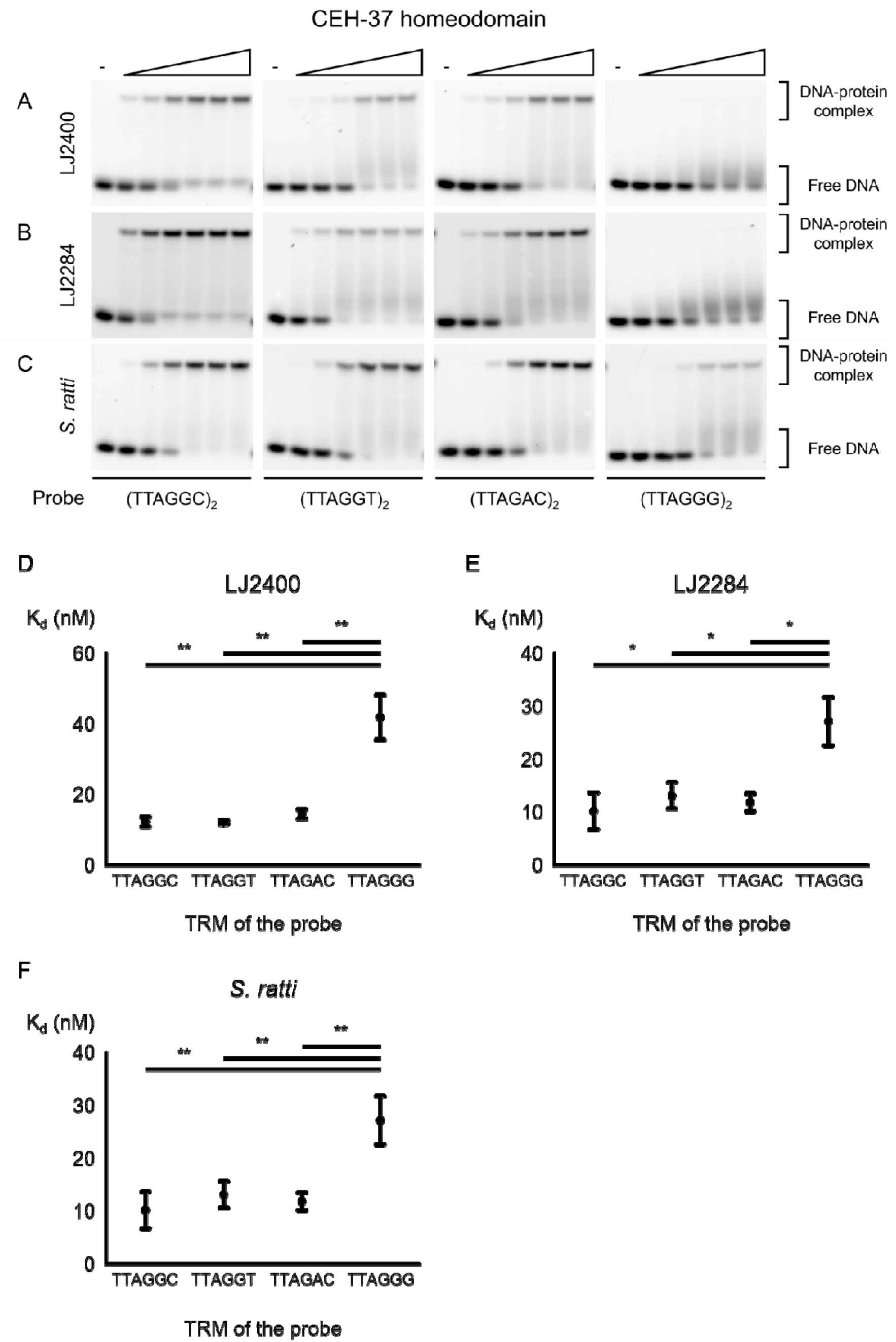
Binding affinity of CEH-37 homeodomains from clade IV nematodes. (A-C) Representative images of EMSA demonstrating the binding of the LJ2400, LJ2284, and *S. ratti* CEH- 37 homeodomains to telomeric DNAs, respectively. 5 nM of DNA probes indicated under each figure were incubated with increasing concentrations of proteins ranging from 5 nM to 160 nM for LJ2284 and *S. ratti* CEH-37 homeodomains besides the concentration range of the LJ2400 protein was 2.8 nM to 90 nM. (D-F) K_d_ of CEH-37 homeodomains from LJ2400, LJ2284, and *S. ratti* binding to each telomeric DNA, respectively. DNA probes used for the K_d_ measurements consist of 2 repeats of the indicated sequences. Relationships without asterisks indicate non-significant differences (Newman-Keuls multiple comparison test, * 0.01<*p*<0.05, ** 0.005<*p*<0.01, *** *p*<0.005).

As our results show that CEH-37 does not effectively bind to telomeric DNA with TTAGGG repeats, we measured the binding affinities of CEH-37 homeodomains from *S. ratti*, the parasitic nematode in Clade IV expected to possess noncanonical TRM TTAGGG (Lim et al., 2023). Interestingly, the CEH-37 homeodomain from *S. ratti* also did not bind efficiently to TTAGGG-repeated DNA compared to other nematode telomeric DNAs (Figure 3C and F). Contrary to a previous study, this result suggests that *S. ratti* may not have TTAGGG as its TRM, or it may rely on a different protein instead of CEH-37.

### Nematodes with novel TRMs need alternative mechanisms to protect their G-rich telomeric overhangs other than capping by POT proteins

TEBP and POT proteins form a telomeric protein complex, which is essential for telomere protection in *C. elegans* (Dietz et al., 2021; Yamamoto et al., 2021). Interestingly, while these proteins are well conserved in *C. uteleia*, a clade V nematode species with distinct TTAGGT TRM, they are not conserved in clade IV nematodes which include other noncanonical TRMs (Figure 1A). To investigate how these proteins contributed to the evolution of TRMs, we obtained *C. elegans* POT proteins, POT- 2 and POT-3, and assessed their binding affinities to nematode telomeric DNAs (Figure 4A and B, Supplementary Figure S4A-C). The EMSA results indicate that both POT-2 and POT-3 bind efficiently to TTAGGC repeats. However, they did not bind to the DNA probe composed of TTAGGT repeats or TTAGAC repeats. These results indicate that POT proteins exhibit high sequence specificity in binding telomeric DNA, failing to bind to the novel nematode telomeric repeats.

**Figure 4.**
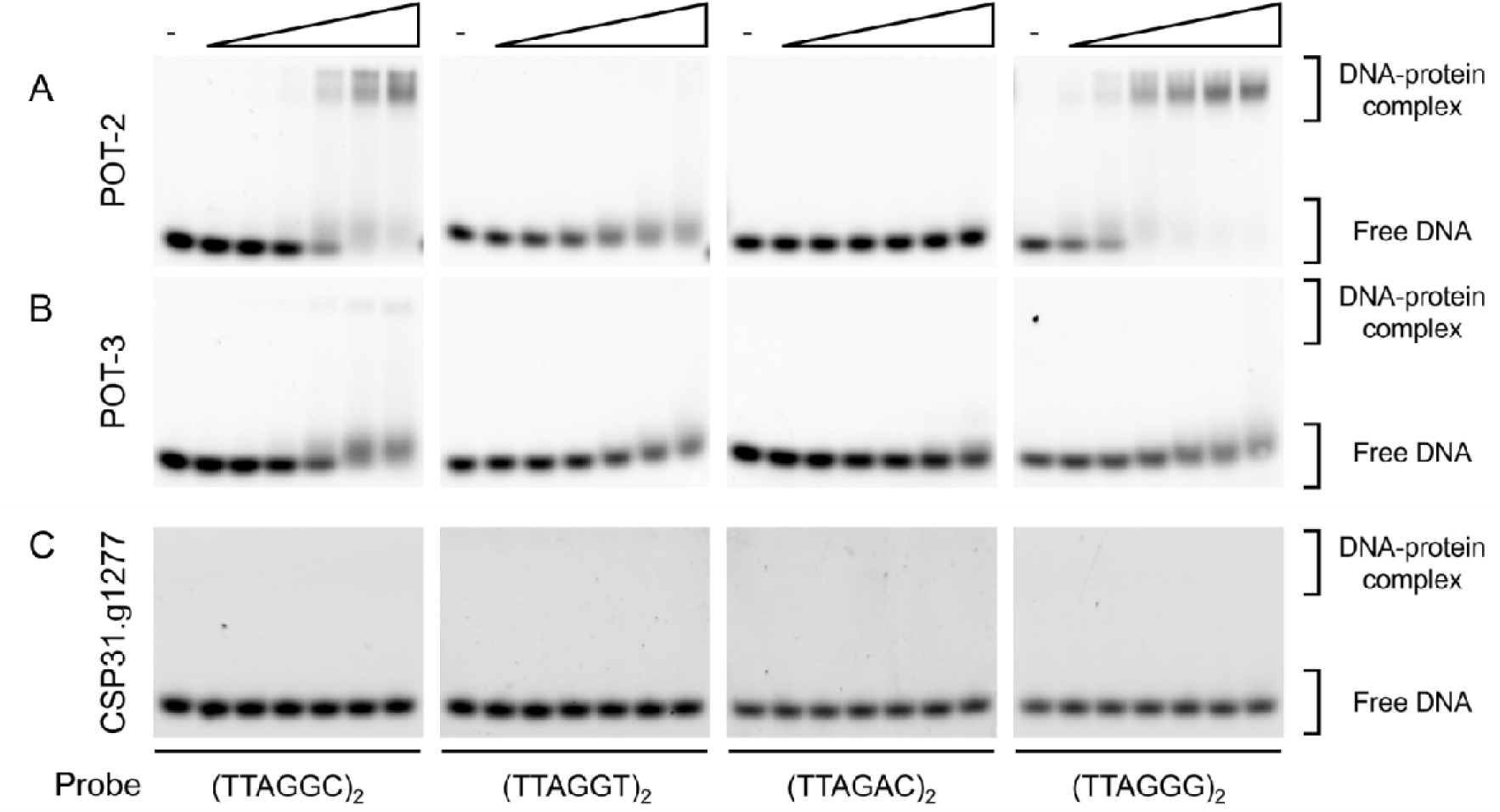
Analysis of binding affinity of recombinant POT-2 and POT-3 to nematode telomeric repeats. (A, B, C) Representative images of EMSA demonstrating the binding of the *C. elegans* POT- 2, POT-3, and *C. uteleia* CSP31.g1277 (a homolog protein of *C. elegans* POT-2 and POT-3) to telomeric DNAs, respectively. 5 nM of DNA probes were incubated with increasing concentrations of proteins ranging from 20 nM to 640 nM except for the reaction by POT-2 and POT-3 to (TTAGGC)_2_ probe, where the protein range was 5 nM to 160 nM.

For the emergence of novel TRMs, POT proteins must either coevolve or be lost. As the results of TBP conservation indicate, POT proteins are not conserved in clade IV nematodes, suggesting that the TTAGAC TRM in clade IV arose by the loss of POT proteins (Figure 1A). To confirm whether this observation is a general phenomenon in clade IV nematodes, we performed a BLASTP search of every *C. elegans* POT protein (POT-1, POT-2, and POT-3) against the protein database of clade IV nematodes, including Panagrolaimidae isolates LJ2284 and LJ2285, which possess the TTAGAC TRM (Supplementary Table S5). As expected, POT proteins were not conserved in any clade IV nematodes, regardless of their TRM. This result suggests that the loss of *pot* genes preceded the emergence of the noncanonical TTAGAC TRM in clade IV. Similar to the earlier case of CEH-37, the loss of POT protein may have reduced selective pressure against the TTAGAC TRM, facilitating the emergence of the noncanonical TRM in Panagrolaimidae isolates.

While the emergence of the TTAGAC TRM can be attributed to the loss of major subunits of the telomeric protein complex, the emergence of the TTAGGT TRM in *C. uteleia* still raises the possibility of coevolution. Unlike clade IV nematodes, BLASTP analysis has shown that *C. uteleia* retains homologs of *C. elegans* POT-1 (CSP31.g24055) and POT-2 and POT-3 (CSP31.g1277). Since CSP31.g1277 (bit score = 137) showed a higher bit score with *C. elegans* POT-2 and POT-3 than CSP31.g24055 (bit score = 73.6) did with POT-1, we purified recombinant CSP31.g1277 to test whether it binds to its host telomeric DNA (Supplementary Figure S4D and E). Surprisingly, CSP31.g1277 failed to bind to any nematode telomeric DNAs (Figure 4C) while it minimally bound regardless to the probe sequence in extreme conditions such as binding reaction at 37°C, overnight (Supplementary Figure S4), indicating that it cannot function as a TBP. Since the binding affinity of CSP31.g24055 to telomeric DNAs has not yet been tested, it remains possible that *C. uteleia* possesses C-rich overhangs where CSP31.g24055 may bind, or that alternative proteins or mechanisms are involved in protecting G-rich telomeric overhangs.

## Discussion

Chromosome ends are protected by the intricate interaction between sequence-specific TBP and telomeric DNA. However, the relationship between the evolution of TRM and TBPs in animals has been elusive. In this study, we examined how TRMs could evolve in the challenge posed by sequence-specificity of TBPs in Nematoda. Phylogenetic analysis and measurements of binding affinities of CEH-37 and POT proteins suggested that the presence of TBPs with its sequence-specificity exert selective forces on TRM evolution.

The evolution of telomeric sequences and their associated proteins has also been observed in yeasts. Yeasts exhibit a wide range of TRMs, from common TTAGGG repeats to long (>20 bp) and heterogeneous TRMs. This extreme diversity accompanied the adoption of flexible TBPs, such as Rap1 and Taz1, instead of the strictly sequence-specific Teb1/Tay1 (Moser and Nakamura, 2009; Sepsiova et al., 2016). As the new TBP emerged (i.e. Taz1) or the telomere-associated protein became a multifunctional TBP by gaining the ability to bind DNA directly (i.e. Rap1), the original TBPs either became dedicated to their secondary role as transcription factors or disappeared. The evolution of telomeres in yeasts, where TBPs have both telomeric functions and roles as transcription factors, is reminiscent of the nematode CEH-37, which is also known for its roles as a transcription factor involved in sensory neuron specification and intestinal immunity in *C. elegans* (Lanjuin et al., 2003; Liu et al., 2023; Shore and Nasmyth, 1987).

Along with these findings, it is plausible that CEH-37 served as a main dsTBP in species lacking TEBP proteins and other *C. elegans* telomeric protein complex subunits. Prior to the discovery of TEBP proteins, CEH-37 was speculated to be a functionally conserved dsTBP based on its structural similarity to Myb-related domains of dsTBPs in other species, such as human TRF1 or *S. pombe* Taz1. TBP conservation analysis in our study supports this view, as CEH-37 is the only TBP with this structural similarity in clades where TEBP proteins are not conserved (Table S2) (Kim et al., 2003; Moon et al., 2014). Therefore, we could cautiously hypothesize that CEH-37 is a result of convergent evolution that may have served as an ancient form of the main dsTBP in nematodes which functions in species lacking TEBP proteins.

The emergence of the TTAGAC TRM in clade IV nematodes can be attributed not only to the flexibility of CEH-37 but also to the absence of specifically binding POT proteins, which hardly bind to TTAGAC repeats. Unlike most eukaryotes, which possess oligonucleotide/oligosaccharide-binding (OB) fold POT proteins to protect their telomeric overhangs, clade IV nematodes lack these POT proteins (Table S5). Clade IV nematodes likely evolved adaptations to compensate for the loss of POT proteins. Various strategies for protecting single-stranded telomeric DNA in the absence of POT proteins have been observed in other organisms. In budding yeast, telomeric overhangs are capped by the Cdc13-Stn1-Ten complex in place of POT proteins (Moser and Nakamura, 2009). In plants, single-stranded telomeric DNA binding proteins (ssTBPs) with or without OB folds have been identified, while some species possess blunt-ended telomeres (Kazda et al., 2012; Luo et al., 2020). These precedents suggest that investigating how nematodes adapt to the loss of POT proteins could be a compelling area for further study, with the potential to discover new mechanisms.

Given that the nematode telomeric protein complex is known to comprise TEBP and POT proteins (Dietz et al., 2021; Yamamoto et al., 2021), it was surprising that a homolog of the 3’ telomeric overhang-binding POT proteins from *C. uteleia*, a unique clade V nematode with a distinct TTAGGT TRM, hardly binds to telomeric DNA. Since the 3’ telomeric overhang-binding POT proteins do not directly associate with the TEBP proteins in *C. elegans*, their functional significance might be reduced in the complex. Consequently, *C. uteleia* may employ other proteins to protect its 3’ telomeric overhangs or lack such overhangs. In contrast, *C. elegans* POT-1, which binds to the 5’ telomeric overhangs, directly interacts with TEBP proteins. Therefore, it is plausible that CSP31.g24055, the *C. uteleia* homolog of POT-1, binds to the novel telomeric DNA. This could have occurred either through the flexible binding specificity of *C. elegans* POT-1 or via the coevolution of CSP31.g24055 within the emergence of the novel TTAGGT TRM. Evidence supporting the former possibility comes from the case of CEH-37 in this study, as well as the flexibility observed in yeast RAP1 and Taz1 (Moser and Nakamura, 2009; Sepsiova et al., 2016). The latter one is supported by studies of plant evolution, where POT1 in green algae only binds to TTTAGGG repeats, while the POT1 of land plants possessing the TTAGGG TRM binds to both TTAGGG and TTTAGGG repeats (Shakirov et al., 2009). Identifying the mechanism by which the *C. uteleia* TRM arose would be an intriguing study.

We confirmed that CEH-37 homeodomains from every species, including *S. ratti*, do not effectively bind to TTAGGG repeats compared to other telomeric DNAs found in Nematoda, raising questions about the nature of the telomere in *S. ratti*. Previous WGS data speculated that *S. ratti* possesses a TTAGGG TRM, as this was the only TTAGGC-related motif identified in the species (Lim et al., 2023). However, TRM-containing concatemers were rare compared to TRM-containing reads from WGS data of other nematodes analyzed in the study, ranking 67^th^ out of 67 species. Based on this information, we hypothesize that 1) *S. ratti* possesses short telomeric DNAs consisting of TTAGGG repeats, possibly protected by TBPs other than CEH-37 or by other mechanisms (e.g. adoption of rigid G-quadruplex (G4) structures), or that 2) *S. ratti* does not possess TTAGGG TRM, and the observed reads from the parasite may have originated from its mammalian host (Bryan, 2020; Smith et al., 2011). If the first hypothesis is true, it would be interesting to investigate the role of G4 structure in telomeres by comparing telomeres of *S. ratti*, nematodes with TTAGGC TRMs which are also known to form telomeric G4 structures, and nematodes with TTAGAC TRMs which are unlikely to form telomeric G4 structures (Marquevielle et al., 2022).

In summary, we demonstrated the dynamics of nematode telomere evolution and highlighted the role of TBPs in the evolution of telomere sequences across different nematode species. We expect our study to not only advance the field of telomere biology using nematodes as multicellular organism models but also enhance our understanding of the evolution of DNA-protein interactions beyond telomeres.

## Experimental procedures

### Strains

Nematodes were grown in nematode growth medium seeded with *Escherichia coli* strain OP50 at 20°C as standard methods (Brenner, 1974). *C. elegans* Bristol N2, *C. uteleia* JU2585, and Panagrolaimidae isolates (LJ2284 and LJ2400) were used for this study.

### Conservation analysis of telomere-binding proteins

FASTA protein files of *C. elegans* (PRJNA13758) and *C. uteleia* (PRJEB12600) were obtained from WormBase ParaSite (Howe et al., 2016; Howe et al., 2017). Protein sequences of Panagrolaimidae isolates (LJ2284, LJ2285, and LJ2400) were kindly produced by Lim et al (Lim et al., 2023).

To analyze conservation of telomere binding proteins of *C. elegans*, we used DIAMOND (version 2.1.8; diamond blastp -d -q -o --threads 20 --very-sensitive and diamond blastp -d -q -o --threads 20 --ultra-sensitive) (Buchfink et al., 2021). The conservation was evaluated by using the highest bit score protein sequence with conserved domains of original proteins identified using NCBI CD-search (CDD database, version 3.21) (Lu et al., 2020; Marchler-Bauer et al., 2017; Wang et al., 2022). Multiple sequence alignment was performed by Clustal Omega using Jalview 2.11.4 (Waterhouse et al., 2009). Additionally, BLAST+ provided by WormBase ParaSite on its web server or DIAMOND as mentioned above were used to analyze conservation of TEBP-POT complex subunits in clade IV nematodes (Camacho et al., 2009). Query sequences were used as written on Table S4 and BLAST was searched against the clade IV nematode protein database.

### Construct generation

*ceh-37* homeobox regions of each strain were cloned in a modified pGEX-4T-1 vector containing a HRV 3C cleavage site after the GST tag while *C. elegans pot-2* and *pot-3* were cloned in a modified pMJ806 vector containing TEV cleavage site after the MBP tag. To generate cDNA of desired genes, RNAs isolated from nematodes with TRIzol were reverse transcripted with PrimeScript RT Master Mix (Takara) except for *S. ratti*. Since we were not able to culture *S. ratti*, *ceh-37* homeobox cDNA of *S. ratti* was generated by ligating two annealed ∼110bp oligos after phosphorylation. cDNAs were fused into the vector by Gibson assembly master mix (New England Biolabs).

### Protein expression and purification

The plasmid containing the desired gene was transformed into *E. coli* Rosetta (DE3) cells. Colonies were picked and inoculated into Terrific Broth, and the culture was grown at 37°C until the OD_600_ reached approximately 0.6. Protein overexpression was induced by adding IPTG to a final concentration of 0.2 mM, followed by incubation at 20°C for 18 h. The cells were harvested and resuspended in PBS, and cell lysis was performed using sonication, supplemented with PMSF and DNase I. The lysed cells were centrifuged at 14,000 rpm for 15 min, and the supernatants were collected. The supernatants from cells expressing CEH-37 or POT proteins were incubated with glutathione agarose resin or amylose agarose resin, respectively, pre-equilibrated with PBS for 1 h with gentle rolling. The resin was washed with 10 column volumes of 20 mM HEPES pH 7.5, 100 mM NaCl buffer. While the proteins were bound to the resin, HRV3C or TEV protease was added for overnight digestion at 4°C. The following day, the protein was released using the wash buffer, and subsequently loaded onto a HiTrap SP (CEH-37)/Q (POT proteins) HP 5 mL column (GE Healthcare). An appropriate salt gradient was applied using 20 mM HEPES pH 7.5 buffer, and peaks were analyzed by SDS-PAGE. The desired fractions were pooled and concentrated. Final purification was performed by loading the concentrated protein onto a Superdex 200 10/300 GL column (GE Healthcare) equilibrated with 20 mM HEPES pH 7.5, 150 mM NaCl. The peaks were analyzed by SDS-PAGE, and the desired fractions were collected and concentrated to the appropriate concentration. The concentrated protein solution was aliquoted, flash-frozen in liquid nitrogen, and stored at -80°C.

During the purification of *C. uteleia* protein CSP31.g1277, the protein precipitated after protease treatment due to the minimal difference between the isoelectric point (7.93) of tag-free CSP31.g1277 and the pH of the buffer (7.5). To address this, we initially purified His-MBP-tagged CSP31.g1277 using a HiTrap Q HP 5 mL column followed by a Superdex 200 10/300 GL column. Subsequently, the buffer was exchanged to 20 mM HEPES pH 7, 150 mM NaCl using dialysis tubing, and His-tagged TEV protease was added for cleavage at 4°C overnight. The tag and protease were removed by Ni-NTA resin (Qiagen), and the supernatant containing purified CSP31.g1277 was collected for further experiments.

### Electrophoretic mobility shift assay (EMSA)

Telomeric DNA substrates used as DNA probes were prepared by annealing oligos with forward strand having Cy5 fluorescent on its 5’ end. 5 nM of DNA probes were incubated with appropriate concentration of proteins in binding buffer (20 mM Tris-HCl, 50 mM NaCl, 1 mM MgCl2, 5 mM DTT, 5% glycerol, 0.001% tween-20, and 50 ug/mL BSA) for 15 min in room temperature. Samples were loaded to 6% native gel that had been pre-run. Electrophoresis was performed for 90 min at 130 V in a cold room with 0.5x TBE as a running buffer.

### Structure prediction and alignment

To predict biomolecular structures, we utilized the AlphaFold Server powered by AlphaFold 3 model (Abramson et al., 2024). Alignments of CEH-37 homeodomains from species with distinct TRMs were performed using the Matchmaker tool in UCSF ChimeraX (version 1.5) (Goddard et al., 2018).

Structural alignments were conducted using the web-based pairwise structure alignment service provided by the RCSB PDB website, employing the TM-align method (Zhang and Skolnick, 2005). The structures of each protein were obtained from models stored in the AlphaFold Protein Structure Database (Jumper et al., 2021; Varadi et al., 2022). The regions of proteins selected for the alignments were pre-determined DNA-binding domains specified in previous studies (König et al., 1996; Lu et al., 2020; Marchler-Bauer et al., 2017; Sigrist et al., 2005; The UniProt Consortium, 2022; Wang et al., 2022).

### Supporting information

This article contains supporting information.

## Acknowledgements

We are grateful to Dr. J. Lim providing the sequencing data of Panagrolaimidae famility isolates (LJ2284, LJ2285, and LJ2400), and Dr. M-A Félix for providing *C. uteleia* JU2585. We also thank the members of Dr. J. Lee (laboratory of genes and development) and Dr. H-J Choi (laboratory of structural biology) for helpful discussion during the study.

## Author contributions

H.S., H.-J.C., and J.L. conceptualization; H.S. data curation; H.S. formal analysis; J.L. funding acquisition; H.S., S.K., and D.S.L. investigation; H.S., S.K., D.S.L., and H.-J.C. methodology; H.-J.C. and J.L. supervision; H.S. visualization; H.S. writing-original draft; H.S., S.K., D.S.L., and J.L. writing-review and editing

## Funding and additional information

This research was supported by the National Research Foundation of Korea (NRF) grant (2020R1A2C3003352) funded by the Korean Ministry of Science and ICT.

## Abbreviations

DBD: DNA-binding domain
dsTBP: double-stranded telomeric DNA binding protein
EMSA: electrophoresis mobility shift assay
G4: G-quadruplex
K_d_: dissociation constant
MCD: Myb-containing domain
OB: oligonucleotide/oligosaccharide-binding
ssTBP: single- stranded telomeric DNA binding protein
TBP: telomere-binding proten
TRM: telomeric repeat motif
WGS: whole genome sequence

## Supplemental Tables

**Table S1.** Homology of *C. elegans* TBP in Nematoda.

**Table S2.** Pairwise structure alignment results of *C. elegans* and other representative species TBPs TM-align.

**Table S3.** ceh-37 coding sequence of each strain used for binding assay.

**Table S4.** Quantification of EMSA results and dissociation constants.

**Table S5.** Results of BLAST analysis of subunits of *C. elegans* TEBP-POT complex to clade IV nematode protein database.

## Supplemental Figures

**Figure S1.**
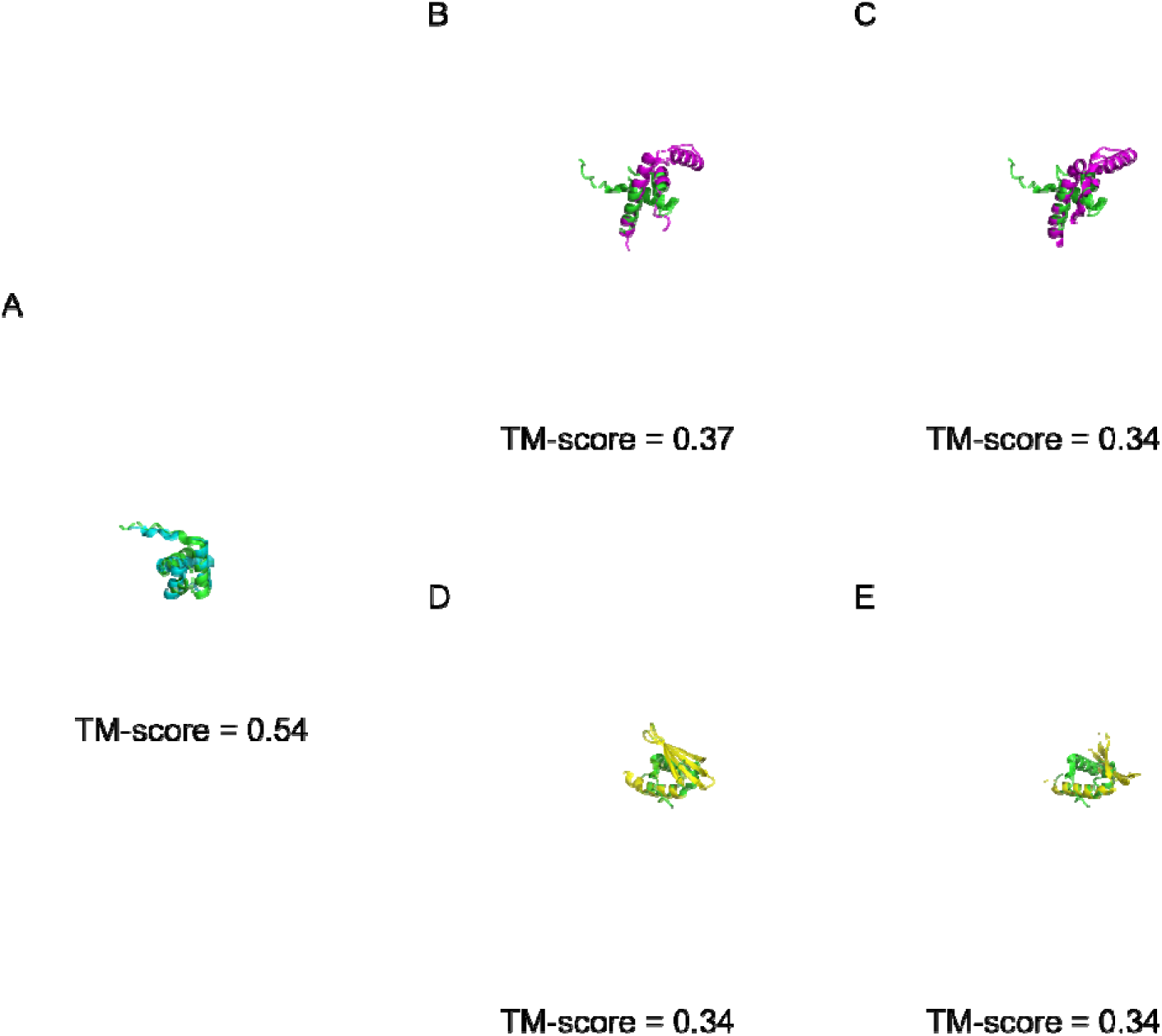
Models of a human TRF1 Myb-like domain aligned to DBDs of *C. elegans* dsTBPs. The Myb-like domain of human TRF1 (green) is structurally aligned to DBDs of (A) CEH-37 (cyan), (B, C) HMG-5 (magenta), and (D, E) PLP-1 (yellow). The unaligned regions of each protein are faintly shown. TM-scores of hTRF1 Myb-like domain to CEH-37 homeodomain, each HMG-box in HMG-5, and each PUR domain in PLP-1 are 0.54, 0.37, 0.34, 0.34, and 0.34, respectively, where TM-score above 0.5 indicates an evolutionary relationship.

**Figure S2.**
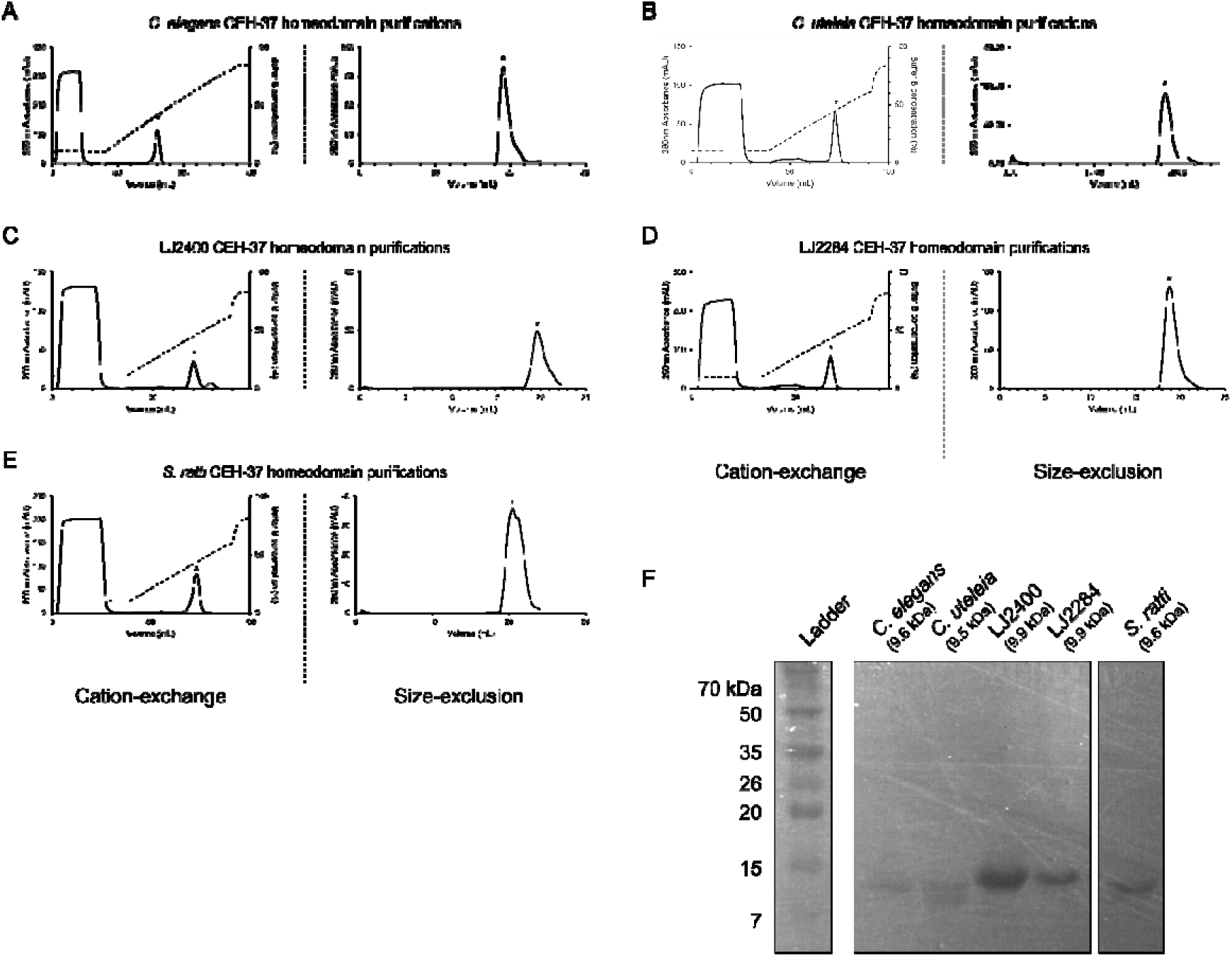
The purification of CEH-37 homeodomains. (A-E) After obtaining CEH-37 homeodomains using a GST-tag in a bacterial expression system, further purification was carried out sequentially using cation-exchange chromatography (left) and size-exclusion chromatography (right). Each graph illustrates the purification results of CEH-37 homeodomains from (A) *C. elegans*, (B) *C. uteleia*, (C) LJ2400, (D) LJ2284, and (E) *S. ratti,* respectively. The asterisks indicate the peaks of CEH-37 homeodomains. (F) The validation of protein purification by SDS-PAGE.

**Figure S3.**
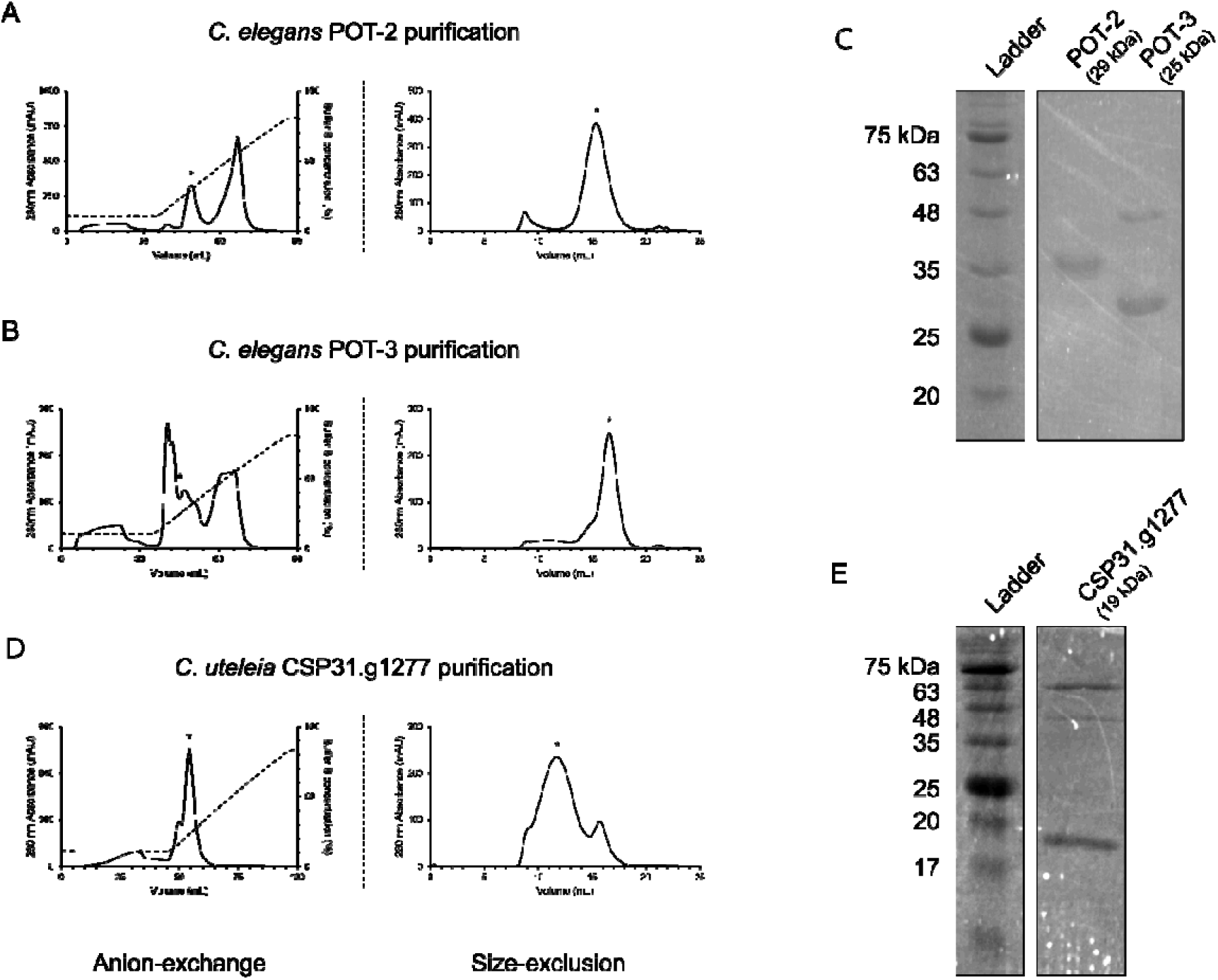
Purifications results of recombinant POT proteins from *C. elegans* and *C. uteleia* obtained by bacterial expression. (A, B) Purification results of *C. elegans* POT-2 and POT-3 by anion-exchange chromatography (left) and size-exclusion chromatography (right), respectively. The asterisks indicate the peak of the target protein. (C) The validation of *C. elegans* POT proteins purification by SDS-PAGE. The upper band on the lane of POT-3 is speculated as a remaining MBP-tag (43 kDa). (D) Purification results of MBP-tagged *C. uteleia* POT protein homolog CSP31.g1277 by anion-exchange chromatography (left) and size-exclusion chromatography (right). (E) The validation of TEV-treated *C. uteleia* CSP31.g1277 purification by SDS-PAGE. The purified sample contained the tagged CSP31.g1277 (62 kDa) and the cleaved tag separated from the protein after TEV treatment (43 kDa).

**Figure S4.**
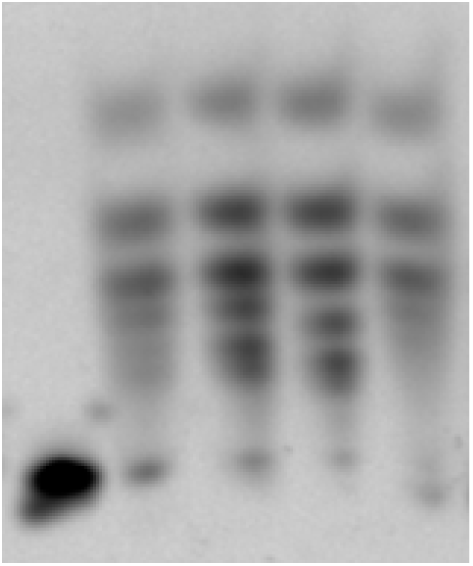
Binding reaction results of the *C. uteleia* POT protein homolog with telomeric DNAs under an extreme condition. DNA probes (5 nM) were incubated with an excess of the protein (>1.7 μM) at 37℃, overnight. This reaction resulted in protein-DNA binding, but the complexes were unstable, which caused multiple bands. The first lane shows the negative control, only containing the (TTAGGC)_2_ DNA probe, while the following lanes contain samples of the protein with (TTAGGC)_2_, (TTAGGT)_2_, (TTAGAC)_2_, and (TTAGGG)_2_ DNA probes in order.

